# Diatoms diversify and turn over faster in freshwater than marine environments

**DOI:** 10.1101/406165

**Authors:** Teofil Nakov, Jeremy Michael Beaulieu, Andrew James Alverson

## Abstract

Many clades that span the marine-freshwater boundary are disproportionately more diverse in the younger, shorter-lived, and scarcer freshwater environments than they are in the marine realm. This disparity is thought to be related to differences in diversification rates between marine and freshwater lineages. However, marine and freshwaters are not ecologically homogeneous, so the study of diversification across the salinity divide should also account for other potentially interacting variables. In diatoms, freshwater and substrate-associated (benthic) lineages are several-fold more diverse than their marine and suspended (planktonic) counterparts. These imbalances provide an excellent system to understand whether these variables interact with diversification. Using multistate hidden-state speciation and extinction models we found that freshwater lineages diversify faster than marine lineages regardless of whether they inhabit the plankton or the benthos. Freshwater lineages also had higher turnover rates (speciation + extinction), suggesting that habitat transitions impact speciation and extinction rates jointly. The plankton-benthos contrast was also consistent with state-dependent diversification, but with modest differences in diversification and turnover rates. Asymmetric, and bidirectional transitions rejected hypotheses about the plankton and freshwaters as absorbing, inescapable habitats. Our results further suggest that the high turnover rate of freshwater diatoms is related to high turnover of freshwater systems themselves.

## Introduction

Freshwater habitats comprise just 2% of the biosphere, yet in lineages that span the salinity gradient the magnitude of freshwater diversity is often comparable, or surpasses, species richness in the marine realm, which covers 70% of Earth’s surface (Mora et al. 2011; Appeltans et al. 2012; Grosberg et al. 2012; Wiens 2015; Miller and Wiens 2017). This is striking given that most of these lineages originated in the oceans, and so—all else being equal—have had more time to accumulate diversity within their ancestral marine environment. Across animals there is a disconnect between species richness and time for accumulation of diversity between marine and freshwaters [e.g., flatworms, rotifers, molluscs, and ray-finned fish (Bloom et al. 2013; Betancur-R et al. 2015; Guinot and Cavin 2015; Wiens 2015; Miller and Wiens 2017)]. Direct tests of asymmetry in diversification have revealed that in amniotes, for example, marine lineages have higher extinction rates (Miller and Wiens 2017; but see Vermeij et al. 2018), whereas marine fish have higher net diversification compared to their freshwater counterparts (Betancur-R et al. 2015). To date, most of the research on this question has focused on animal systems, where there does not appear to be a clear consensus on what is driving disparities in species richness across the marine-freshwater divide (Betancur-R et al. 2015; Guinot and Cavin 2015; Wiens 2015; Miller and Wiens 2017). Microbes, including protists, might provide novel insights into these questions, as they are diverse at both ends of the salinity gradient (Mora et al. 2011; Appeltans et al. 2012) and offer alternative perspectives due to their vastly different dispersal capabilities, population sizes, and physiological features compared to animals.

Here we focus on diatoms, a species-rich lineage of unicellular eukaryotic algae that play important roles in the global cycling of carbon and oxygen and form the base of many aquatic food webs (Armbrust 2009). Although, our knowledge of diatom diversity is far from complete, a clear pattern of imbalanced species richness across the marine-freshwater divide exists, with many more species known from freshwaters than the ancestral marine environment (Kociolek et al. 2017). Most diatom genera are thought to be restricted to either marine or freshwaters (Round and Sims 1980; Round et al. 1990; Mann 1999), and transitions between these environments are thought to be challenging, as migrants need to mitigate osmotic stress linked to changes in cellular water potential, salt stress caused by increased uptake or loss of ions, and changes in the ratios of intracellular ions (Kirst 1990, 1996). These considerations form the basis of a long-held view that colonizations of freshwaters by diatoms are infrequent and irreversible events (e.g., the Rubicon hypothesis of Mann 1999). This hypothesis posits that the modern freshwater flora has been assembled from a small set of phylogenetically diverse colonizers (rarity) and that once adapted to freshwaters, these lineages rarely, if ever, cross back to their ancestral marine habitat (irreversibility), implying that freshwaters are an ‘absorbing’ state. Although the freshwater diatom flora is clearly a composite of many lineages that trace back to independent colonization events (Alverson et al. 2007; Ruck et al. 2016), the frequency of colonizations and irreversibility aspect of the Rubicon hypothesis have only been studied in small clades, making it difficult to derive broad conclusions (Beaulieu and O’Meara 2018). Moreover, studies to date have not explored the interaction between habitat transitions and rates of diversification, which is essential for tests of character-dependent diversification (Maddison 2006).

An important aspect that studies of diversification in marine and freshwaters have yet to consider is that the focal contrast (marine vs. freshwater) is almost certainly not the sole determinant of diversification dynamics, but rather, observed patterns are likely the product of many potentially interacting variables related to the habitats and traits of different lineages (Beaulieu and O’Meara 2016; Caetano et al. 2018). In protists, and in diatoms in particular, the plankton (suspended growth habit) and benthos (substrate-associated growth habit) represent another prominent environmental contrast, with an imbalanced species richness similar in magnitude to the marine-freshwater divide. In diatoms, transitions between planktonic and benthic habitats appear to be linked to both growth habit (e.g., colonial vs. solitary, Nakov et al. 2015) and mode of sexual reproduction (e.g., oogamous vs. anisogamous, Kooistra et al. 2007, 2009). For example, the numerous, flagellum-propelled male gametes of oogamous species might facilitate colonization of the plankton, whereas the few, aflagellated gametes of isogamous lineages might hinder survival and reproduction in dilute planktonic environments (Kooistra et al. 2007, 2009).

Several additional lines of evidence point to potential interactions between the marine-freshwater and plankton-benthos divides. First, the age and longevity of most freshwater lakes supporting planktonic communities are linked to Earth’s glaciation cycles, with only a few dozen long-lived tectonic basins whose formation predates the last interglacial (ca. 130-115 thousand years ago; Mackay et al. 2010; Hampton et al. 2018). The majority of freshwater lakes are geologically recent and have relatively short lifespans, so freshwater planktonic lineages might turn over faster relative to the marine planktonic lineages from environments that have existed continuously since the formation of the oceans. This hypothesis is supported by observed changes the composition of planktonic diatom communities in sedimentary records from ancient lakes that closely follow glaciation and drought cycles (Edlund and Stoermer 2000; Cohen et al. 2007; Mackay et al. 2010; Stone et al. 2011).

Marine and freshwater plankton might also differ in population size and connectivity. Marine populations have fewer physical barriers to migration and are generally expected to have broader geographic distributions and higher rates of gene flow (Palumbi 1992, 1994). As a result, marine planktonic diatoms might experience lower speciation and extinction rates compared to their freshwater counterparts, which inhabit discontinuous and irregularly distributed habitats. Still, in diatoms, gene flow can be limited among marine populations due to isolation by distance (Casteleyn et al. 2010) or ecology (Whittaker and Rynearson 2017), sometimes even at small spatial scales (Rynearson and Armbrust 2004; Godhe and Härnström 2010; Sjöqvist et al. 2015). These findings suggest that population subdivision might be even stronger in freshwaters (Vanormelingen et al. 2008; Evans et al. 2009). Taken together, these observations suggest that observed differences in diversification dynamics across the marine-freshwater divide are confounded by differences in, for example, lineages inhabiting the marine plankton vs. freshwater plankton or the marine plankton vs. marine benthos.

Here, we characterize diatom evolution across the marine-freshwater and plankton-benthos divides using a large, time-calibrated phylogeny representing the known breadth of extant diatom diversity. To account for variation in rates of diversification and colonizations across environments, we used a hidden state speciation and extinction modeling framework (HiSSE; Beaulieu and O’Meara 2016) modified for multiple characters. In doing so, we were able to jointly estimate rates of colonization and diversification across multiple environments and account for possible interactions between focal characters (i.e., marine plankton, marine benthos, freshwater plankton, and freshwater benthos). We used this framework to test two sets of hypotheses. First, we assessed whether diatom diversification (speciation - extinction) or turnover (speciation + extinction) is faster in freshwaters due to, for example, differences in habitat longevity or population size and connectivity. We test our marine-freshwater hypotheses in two ways: irrespective of whether they have a suspended (plankton) or substrate-associated (benthos) habit as well as by distinguishing between marine vs. freshwater plankton and marine vs. freshwater benthos. Second, we tested several hypotheses about the directionality and reversibility of transitions between habitats, given the traditionally held view that both plankton and freshwaters are ‘absorbing’ environments where colonists invade, adapt, and specialize to the point that emigration is unlikely or impossible. We discuss the implications of our findings for diatom ancestry, diversification, and phylogenetic diversity across the ecologically important marine-freshwater and plankton-benthos divides.

## Materials and Methods

### Phylogeny and diversity data

We used a time-calibrated phylogeny of 1,151 diatoms reconstructed from a concatenated alignment of 11 genes (Nakov et al. 2018)—the largest available phylogenetic tree of diatoms that covers their known phylogenetic and ecological breadth. We scored 1,132 out of the 1,151 species for two characters: "habitat" (plankton or benthos) and "salinity" (marine or freshwater) based on taxonomic compilations (Round et al. 1990; Hasle and Syvertsen 1996; Lange-Bertalot and Krammer 2000; Witkowski et al. 2000; Lobban et al. 2012) and previous work on character evolution (Alverson et al. 2007; Nakov et al. 2015; Ruck et al. 2016).

Given a total estimated diatom species richness of up to 100,000 species (Mann and Vanormelingen 2013), this phylogeny, although the most taxonomically inclusive to date, still falls well short of the full diversity of diatoms. To account for under-sampling in our diversification analyses, we estimated sampling fractions for each environment, e.g., the fraction of sampled marine species from the total number of marine species described to date. Data for salinity (marine-freshwater) were obtained from DiatomBase (Kociolek et al. 2017), which reports the total estimated number of species per genus and the total estimated number of marine species per genus, allowing for a straightforward calculation of the number of non-marine (freshwater) species. Similar data were not available for habitat (plankton vs. benthos), so we used the primary taxonomic literature (Round et al. 1990; Hasle and Syvertsen 1996; Lange-Bertalot and Krammer 2000; Witkowski et al. 2000; Lobban et al. 2012) to record the number of species per genus occurring in the plankton or benthos. Based on these numbers, we then derived a per-genus fraction of planktonic species and multiplied the total estimated genus-level diversity by this fraction to obtain an estimate of the number of planktonic and benthic species per genus. We summed these numbers across all genera represented on our tree to make diatom-wide estimates of species richness in each environment.

### Analysis of the binary salinity and habitat characters

Hypotheses about associations between lineage diversification and discretely coded traits or environments can be tested using state speciation and extinction models (-SSE; Maddison et al. 2007; FitzJohn et al. 2009; Goldberg et al. 2011; Beaulieu and O’Meara 2016). These models jointly estimate rates of evolution of a focal character (i.e., transitions between states) and rates of diversification (i.e., speciation and extinction), thereby disentangling whether an imbalanced ratio of diversity between states reflects differences in diversification or asymmetry in rates of colonization (Maddison 2006). SSE models come in two flavors, ones that assume the rates of speciation, extinction, and transition are shared by all lineages occupying a particular state (e.g., BiSSE, MuSSE, GeoSSE) and those that relax this assumption through the usage of "hidden states" (HiSSE, HiGeoSSE). A third set of models, called character-independent models (CID), allow rate variation independent of the focal characters, and serve as non-trivial null models for state-dependent diversification (Beaulieu and O’Meara 2016; Caetano et al. 2018).

Initially, for each binary character we fit a total of five models: **(1)** a "trivial null" model (number of parameters, k=3, Fig. S1) with one parameter for the turnover rate (defined as speciation + extinction, which measures the rate of any event, birth or death, per unit time) for both observed states, and one parameter controlling the rate of character transitions (e.g., marine <-> freshwater); **(2)** a BiSSE model (k=5, Fig. S1) with separate turnover rates estimated for lineages living in alternate environments and asymmetric transitions between environments (e.g., marine -> freshwater ≠ freshwater -> marine); **(3)** a HiSSE model (k=13) with separate turnover rates for each combination of observed and hidden states and eight transition rates for changes between states (e.g., marine hidden state A -> marine hidden state B). This model disallowed simultaneous transitions of the observed and hidden states (e.g., the rate marine hidden state A -> freshwater hidden state B was set to zero; Fig. S1); **(4)** a CID2 model (k=6) with two turnover rate parameters similar to the BiSSE model, but not linked to the observed character states; and **(5)** a CID4 model (k=8) with five diversification parameters matching the complexity of HiSSE (Fig. S1). In all cases, we kept the extinction fraction (defined as extinction/ speciation, which measures the ratio of death and birth events per unit time) constant across states.

### Analysis of a combined habitat + salinity characters

For the combined habitat + salinity character, we extended the HiSSE and CID framework to handle a four-state case [i.e., marine plankton (MP), marine benthos (MB), freshwater plankton (FP), and freshwater benthos (FB)]. For this analysis, we again fit a trivial null model that assumed no variation in diversification as well as a multistate speciation and extinction model (MuSSE; FitzJohn et al. 2009) that linked diversification rates to the four character states but assumed no variation within regimes (Fig. S1). We then fit two multistate speciation and extinction models with hidden traits, MuHiSSE, and seven multistate character-independent models (MuCID) with increasing numbers of hidden states (MuCID2 through MuCID8; Fig S1). The most complex MuCID8 model had eight parameters for turnover and one for the extinction fraction, matching the complexity of the MuHiSSE model but with diversification parameters not linked to the observed character states (Fig. S1). For both types of models, we allowed transition rates between environments to differ (e.g., MB -> MP ≠ MP -> MB), and we again disallowed dual transitions (e.g., MB -> FP = 0). The two MuHiSSE models differed in how they handled transitions between hidden states. The first model (MuHiSSE reduced) assumed one rate for all transitions between hidden states (e.g., MB hidden state A -> MB hidden state B = FP hidden state A -> FB hidden state B), while the second model (MuHiSSE full) allowed separate transition rates between hidden states (e.g., MB hidden state A -> MB hidden state B ≠ MB hidden state B -> MB hidden state A; Fig S1). Finally, we assessed two alternative implementations of the conditioning for survival of crown lineages, including the original diversitree and HiSSE implementations (FitzJohn et al. 2009; Beaulieu and O’Meara 2016) (HiSSE setting root.type="madfitz") and the conditioning scheme introduced by Herrera-Alsina et al. (2019) (HiSSE setting root.type="herr_als").

In all cases, the extinction fraction was fixed across observed and hidden states regardless of whether we allowed separate estimates of turnover rate for each state.. It is important to note, however, that although extinction fraction is constrained among states, converting turnover and extinction fraction into speciation and extinction allows estimation of separate birth and death rates for each state (Beaulieu and O’Meara 2016). We accounted for incomplete sampling by incorporating the sampling fractions described above directly into our diversification analyses (FitzJohn et al. 2009; Beaulieu and O’Meara 2016). This approach assumed random sampling of each observed character state. For all models, we aimed for 50 maximum likelihood optimizations, each initiated from a different starting point.

### Diversification and transition hypotheses

After obtaining initial results on the fits of state-dependent and state-independent models, we tested a second set of state-dependent models that constrained certain parameters to equality or to zero, effectively removing them from the model. These models were designed to test specific hypotheses related to asymmetry in rates of species turnover (speciation + extinction) or transitions between environments. For these models, we constrained or dropped parameters for each hidden state, permitting rate heterogeneity in the presence of our constraints and maintaining a level of complexity comparable to MuHiSSE. For example, when testing whether the plankton in freshwaters is an absorbing, dead-end state from which lineages rarely or never emigrate, we fixed the plankton-to-benthos transition to zero in both hidden states but still allowed variation in diversification rates within the freshwater plankton and in the rate of colonization of the marine plankton from the freshwater plankton.

To assess the association between the plankton—benthos contrast and diversification, we forced the turnover rate to be the same between: **(1)** the plankton and benthos in marine environments (MP = MB), **(2)** the plankton and benthos in freshwater environments (FP = FB), and **(3)** the previous two combined (MP = MB and FP = FB). These three models tested specifically for differences in diversification dynamics related to suspended vs. substrate-associated growth habit at either or both ends of the salinity gradient. We also evaluated whether the marine-freshwater contrast mattered in the context of one type of growth habitat, but not the other. For example, it could be that benthic lineages differ in turnover between marine and fresh waters, whereas this distinction is less important in the plankton. This set of hypotheses included models that forced turnover to be the same between: **(4)** the marine and freshwater plankton (MP = FP), **(5)** marine and freshwater benthos (MB = FB), and **(6)** the two previous combined (MP = FP and MB = FB).

We also tested hypotheses related to the directionality of transitions between habitats, given the historical understanding of freshwaters and the plankton as absorbing states. In modeling terms, these hypotheses translate to strongly asymmetric transition rates, with the rates of transition outside of the putatively absorbing state equal to zero. For freshwaters, we set up models in which: **(7)** the freshwater plankton is absorbing (FP -> MP = 0, but FP -> FB ≠ 0), **(8)** the freshwater benthos is absorbing (FB -> MB = 0, but FB -> FP ≠ 0) and **(9)** freshwaters in general are absorbing irrespective of growth habit (FP -> MP = 0 *and* FB -> MB = 0). For the plankton, we had an analogous setup with models in which **(10)** freshwater plankton is absorbing (FP -> FB = 0, but FP -> MP ≠ 0), **(11)** the marine plankton is absorbing (MP -> MB = 0, but MP -> FP ≠ 0) and **(12)** plankton is absorbing in both marine and freshwaters (FP -> FB = 0 *and* MP -> MB = 0). In all cases, we disabled only the transitions reflecting the change in state in the trait of interest while allowing transitions between states of the other trait. For example, for freshwater plankton, when we tested whether freshwaters were absorbing, we disallowed the transition to marine plankton and allowed migration to the freshwater benthos, but when we tested whether the plankton was absorbing, we disallowed the transition to the freshwater benthos and allowed colonization of the marine plankton.

### Post-processing of HiSSE and MuHiSSE model fits

In addition to estimating the most likely parameter values for predefined regimes, the models included in HiSSE also allowed us to reconstruct the probabilities of the hidden states occupied by each lineage. For example, a lineage might be freshwater, but its hidden state might be less certain, with some probability of occupying either of the hidden states in the model. In such cases it is unclear which parameter estimate should be assigned to a particular lineage because the diversification and transition rates might differ substantially among hidden states. One way to account for this uncertainty is to average the parameter estimates weighted by the probabilities of occupying alternative hidden states. For example, if the model has two hidden marine states with turnover waiting times of 5 and 10 million years respectively, and a species was reconstructed as occupying marine hidden state *A* with probability of 0.3 and marine hidden state *B* with probability of 0.7, then the waiting time assigned to it, reflecting the uncertainty in the ancestral state reconstruction, would be the weighted average 5*0.3 + 10*0.7 = 8.5. The alternative would be to disregard the uncertainty in the reconstruction and assume that the species in question simply occupies marine hidden state *B* (the more likely state) and assign to it longer waiting time of 10 million years. This type of averaging of per-lineage diversification parameters provides a measure of rate variation within regimes, because instead of splitting species and their rates within a regime into two groups corresponding to the two hidden states, we can obtain a distribution of weighted averages of the parameters. Even if character-dependent diversification is supported, it is unlikely that all lineages assigned to a regime would have diversified at the same rate, and this variation will be reflected in the hidden state reconstructions. This model-averaging approach also alleviates the subjectivity of choosing arbitrary thresholds for choosing models, and instead puts the emphasis of interpretation on the parameter estimates (see Caetano et al., 2018 for further discussion of this point).

We reconstructed marginal ancestral states either from a single best model, when support measured by the Akaike weight was > 99% or from all models supported by ≥ 1% (HiSSE functions MarginRecon and MarginReconMuHiSSE). A vignette with further details regarding MuHiSSE is available as part of the hisse R package. The data and scripts used here are available from a Zenodo repository (DOI: 10.5281/zenodo.2558448). Model summaries and figures for HiSSE objects can also be made with our R package "utilhisse" (https://github.com/teofiln/utilhisse) and an interactive web application (https://diatom.shinyapps.io/hisse-web).

We also approximated the likelihood surface around the maximum likelihood estimates using HiSSE’s adaptive sampling routine (HiSSE functions SupportRegion and SupportRegionMuHiSSE, Beaulieu & O’Meara 2016). This procedure samples the parameter space around the MLEs and re-estimates the likelihood to find a range of parameter values whose likelihood falls within 2 log-likelihood units of the best fit. A broad range would indicate a flatter likelihood surface and less confidence in the parameter estimates. How far the sampler wanders away from an MLE can be controlled with a scaling argument.

## Results

### Binary characters

Diatom diversity data showed strong disparities in species richness across both the marine-freshwater and plankton-benthos divides (Mora et al. 2011; Kociolek et al. 2017). Approximately 70% of the species described to date are freshwater and 83% are benthic. The estimated numbers of species in marine and freshwater habitats are 3221 and 7349 species respectively, and of these our phylogeny includes 680 (21%) marine and 452 (6%) freshwater species. The estimated numbers of species in plankton and benthos are 1800 and 8770 species respectively, and of these our phylogeny includes 358 (20%) planktonic and 774 (9%) benthic species.

These imbalanced diversity ratios suggest that diversification rates might be related to one or both of these characters. Indeed, BiSSE models that linked diversification and character transition rates to the alternate environments provided a much better fit than models that assumed one rate across the tree (Table 1). Freshwater and benthic lineages, as expected given the diversity ratios, had waiting times (inverse of the rate) of speciation + extinction (i.e., net turnover) that were 3.8 and 1.6 times faster, respectively, than their marine and planktonic counterparts. Similarly, net diversification (speciation - extinction) was 3.7 times faster in freshwater than marine lineages, and 1.6 times faster in benthic than planktonic diatoms. The ratio of extinction to speciation (the extinction fraction) was high in both cases: 0.86 for the marine-freshwater and 0.87 for the plankton-benthos contrast. According to BiSSE models, both of these environmental gradients appear to have been important in shaping the diversity of diatoms.

**Table 1.** Comparison of character-dependent and character-independent models for the binary coding of salinity and habitat. The models favored by AICc are in bold. Abbreviations: npd, number of diversification parameters (turnover + extinction fraction); npt, number of transition parameters; Δ, difference in AICc; ω, Akaike weight; F, freshwater; P, plankton.

| Model | lnL | npd | npt | AICc | $\Delta$ | $\omega$ |
| --- | --- | --- | --- | --- | --- | --- |
| Salinity (Marine vs. Freshwater) |  |  |  |  |  |  |
| **hisse** | -5325.72 | **5** | **8** | 10677.77 | 0 | **0.999** |
| cid4 | -5341.02 | 5 | 3 | 10698.16 | 20.39 | <0.001 |
| hisse F absorbing | -5393.10 | 5 | 6 | 10808.44 | 130.67 | <0.001 |
| cid2 | -5444.22 | 3 | 3 | 10900.52 | 222.75 | <0.001 |
| bisse | -5506.97 | 3 | 2 | 11023.99 | 346.22 | <0.001 |
| Habitat (Plankton vs. Benthos) |  |  |  |  |  |  |
| **hisse** | -5164.73 | **5** | **8** | 10355.78 | 0 | **0.999** |
| cid4 (three rates) | -5189.49 | 5 | 3 | 10395.11 | 39.33 | <0.001 |
| hisse P absorbing | -5189.70 | 5 | 6 | 10401.63 | 45.85 | <0.001 |
| cid2 | -5250.83 | 3 | 3 | 10513.74 | 157.96 | <0.001 |
| bisse | -5377.10 | 3 | 2 | 10764.25 | 408.46 | <0.001 |

To assess variation in diversification and transition rates within regimes, we also evaluated character-dependent (HiSSE) and character-independent (CID) models that included hidden states, i.e., factors other than our focal characters that might have influenced diversification. For both characters, we found substantial variation in the rates of diversification and transitions between regimes, as inferred from the stronger support for models with hidden states relative to BiSSE models (Table 1). A character-dependent HiSSE model was favored for both binary traits with Akaike weights of **ω**=0.999 (Table 1). Generally, freshwater lineages speciated, but also went extinct, faster than marine lineages, resulting in faster overall species turnover in freshwater (Table 2), but with variation among hidden states. The difference between speciation and extinction (net diversification) favored marine lineages in one hidden state (rate ratio marine:freshwater = 2.8), and freshwater lineages in the other hidden state (rate ratio freshwater:marine = 13.4; Table 2). For habitat, the differences in waiting times to a net gain of one species between planktonic and benthic lineages were less pronounced, with plankton:benthos rate ratios ranging from 1.05 in one hidden state to 1.2 in the other (Table 3).

**Table 2.** Parameter estimates (events per million years) and 2 log-likelihood unit confidence interval (in brackets) for the best model (hidden-state speciation and extinction, HiSSE) for salinity (marine vs. freshwater). The letters next to character states indicate the hidden states.

|  | Marine A | Freshwater A | Marine B | Freshwater B |
| --- | --- | --- | --- | --- |
| **Transition rates** |  |  |  |  |
| Marine A | - | 0.0150<br>[0.0141-0.0166] | 0.0459<br>[0.0424-0.0509] | 0 |
| Freshwater A | 0<br>[0.0-0.0009] | - | 0 | 0.0187<br>[0.0142-0.0203] |
| Marine B | 0.0029<br>[0.0026-0.0033] | 0 | - | 0<br>[0.0-0.0005] |
| Freshwater B | 0 | 0.1814<br>[0.162-0.196] | 0.0067<br>[0.0050-0.0068] | - |
| **Diversification rates** |  |  |  |  |
| Speciation | 0.168 [0.165-0.183] | 0.059 [0.056-0.065] | 0.035 [0.035-0.04] | 0.472 [0.445-0.52] |
| Extinction | 0.082 [0.08-0.091] | 0.029 [0.028-0.033] | 0.017 [0.017-0.02] | 0.232 [0.221-0.262] |
| Net turnover<br>(speciation + extinction) | 0.25 [0.247-0.274] | 0.089 [0.085-0.098] | 0.053 [0.053-0.06] | 0.704 [0.667-0.772] |
| Net diversification<br>(speciation - extinction) | 0.085 [0.082-0.092] | 0.03 [0.027-0.034] | 0.018 [0.017-0.02] | 0.241 [0.222-0.268] |
| Extinction fraction<br>(extinction/speciation) | 0.49 [0.47-0.519] |  |  |  |

**Table 3.** Parameter estimates (events per million years) and 2 log-likelihood unit confidence interval (in brackets) for the best model (hidden-state speciation and extinction, HiSSE) for habitat (plankton vs. benthos). The letters next to character states indicate the hidden states.

|  | Plankton A | Benthos A | Plankton B | Benthos B |
| --- | --- | --- | --- | --- |
| **Transition rates** |  |  |  |  |
| Plankton A | - | 0.0017 [0.0015-0.0019] | 0.0027 [0.0012-0.0051] | 0 |
| Benthos A | 0.0003 [0.0002-0.0007] | - | 0 | 0.0022 [0.002-0.0023] |
| Plankton B | 0.0955 [0.0842-0.0981] | 0 | - | 0.007 [0.0065-0.0073] |
| Benthos B | 0 | 0.0497 [0.049-0.0608] | 0.0013 [0.0011-0.0016] | - |
| **Diversification rates** |  |  |  |  |
| Speciation | 0.078 [0.065-0.087] | 0.082 [0.075-0.095] | 0.581 [0.521-0.608] | 0.481 [0.449-0.515] |
| Extinction | 0.064 [0.053-0.072] | 0.068 [0.061-0.079] | 0.479 [0.418-0.505] | 0.396 [0.36-0.423] |
| Net turnover<br>(speciation + extinction) | 0.143 [0.118-0.159] | 0.15 [0.136-0.173] | 1.06 [0.939-1.112] | 0.878 [0.809-0.938] |
| Net diversification<br>(speciation - extinction) | 0.014 [0.011-0.016] | 0.015 [0.013-0.019] | 0.103 [0.092-0.116] | 0.085 [0.081-0.097] |
| Extinction fraction<br>(extinction/speciation) | 0.824 [0.8-0.836] |  |  |  |

An alternative way to assess these environmental contrasts is to compare the distributions of diversification rates after accounting for uncertainty in ancestral reconstructions. Waiting time to a turnover event [either speciation or extinction in My calculated as 1/(speciation+extinction)] in marine lineages was on average 11.5 My with a standard deviation (SD) of 4.2 My, whereas freshwater diatoms had a mean waiting time of 3.9 My and SD = 1.4 My between turnover events (Fig. 1A). In contrast, although a HiSSE model was favored, the ranges for turnover waiting time in planktonic (4.5 ± 1.8 My) and benthic (3.28 ± 1.6 My) species were overlapping (Fig. 1B). The variation in net diversification was proportional to turnover (Fig. 1A,B) and for both binary characters these tests highlighted broad rate variation in diversification rates within regimes (Fig. 1A,B).

**Figure 1.**
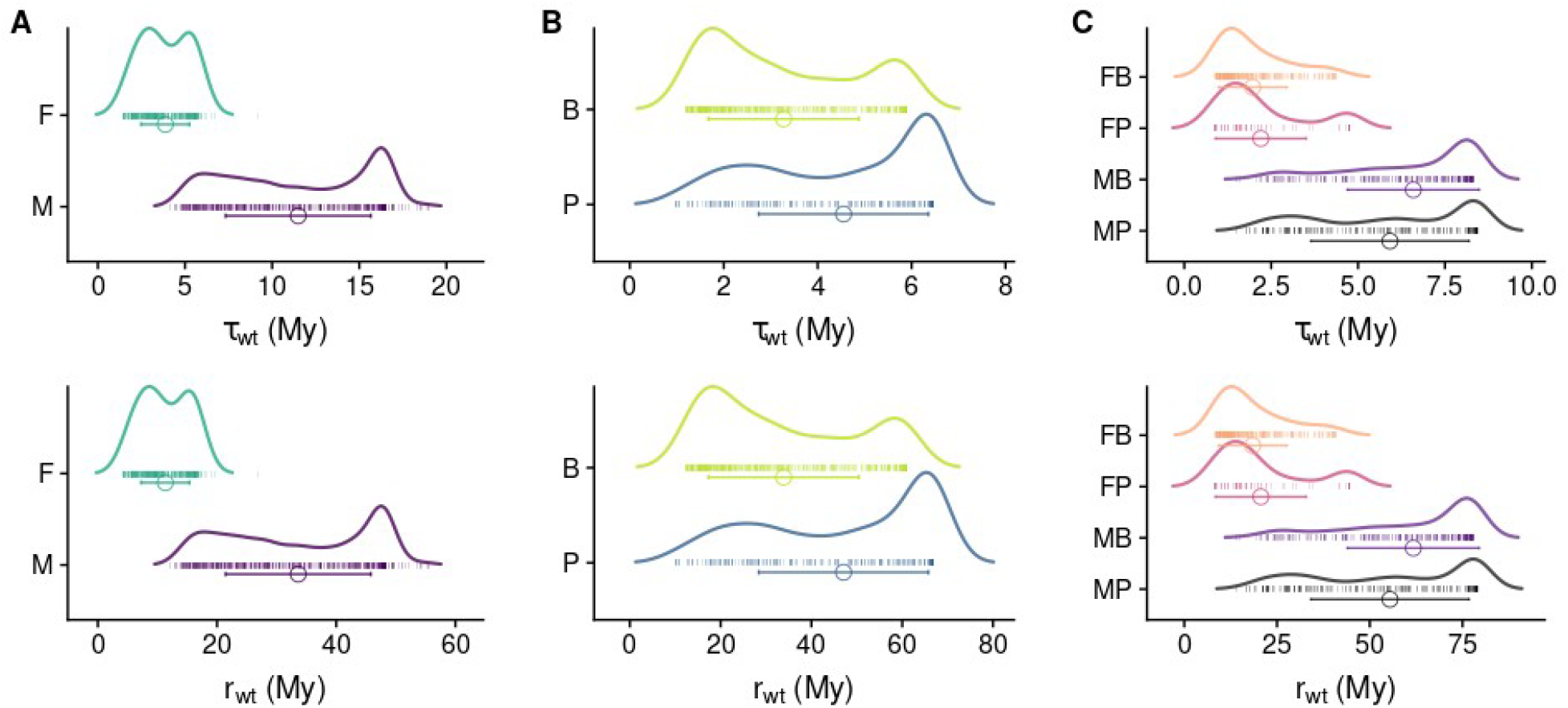
Distributions of tip estimates of net turnover (*τ*, speciation + extinction) and net diversification (***r***, speciation - extinction) averaged across hidden states. (A) binary coding for marine vs. freshwater (HiSSE model), (B) binary coding for planktonic vs. benthic (CID4 model) and (C) multistate coding for the combination of salinity and habitat [averaged across four MuHiSSE variants (Table 1)]. The circles and error bars show the mean ± standard deviation across tips of the phylogeny and the carpet points show the individual rates. The rates are inverted into waiting times (wt) in millions of years. Abbreviations: M, marine; F, freshwater; P, plankton; B, benthos.

We tested hypotheses regarding the directionality of transitions between environments by disabling the transition rates plankton -> benthos and freshwater -> marine (to enforce freshwaters and planktonic environments to be absorbing). In both cases, these models were poorly supported, suggesting that transitions across both environmental gradients are bidirectional (’hisse F absorbing’ and hisse P absorbing’; Table 1).

### Combined salinity + habitat character

One way to further evaluate rate-variation within states is to combine salinity and habitat into a single four-state character: marine-plankton, marine-benthos, freshwater-plankton, and freshwater-benthos. With this approach, we can specifically test for differences between, for example, plankton and benthos, while allowing for the possibility that planktonic lineages in fresh vs. saline waters also differ in diversification. To assess diversification rate differences across the four habitat + salinity regimes, we implemented a multistate variant of the HiSSE model (MuHiSSE), with a separate turnover parameter for each of the four states and two hidden states within each character combination. To guard against false-positive selection of these highly parameterized models, we designed and tested a set of multistate character-independent (MuCID) models with increasing numbers of diversification parameters, up to a MuCID model with eight hidden traits matching the complexity of the MuHiSSE model [eight turnover and one extinction fraction that can be transformed into eight speciation and eight extinction rates (Beaulieu and O’Meara 2016; HiSSE functions ParameterTransform or ParameterTransformMuHiSSE, Fig. S1).

Data for the combined salinity + habitat states mirrored disparities in species richness observed for the individual binary traits. Marine plankton accounted for 11% (1155 species), marine benthos 20% (2066 species), freshwater plankton 6% (645), and freshwater benthos 63% of all diatom species (6704). Our phylogeny included 288 (14%) marine planktonic species, 380 (18%) marine benthic species, 69 (11%) freshwater planktonic species, and 381 (6%) freshwater benthic species.

Marine planktonic diatoms generally have much deeper fossil records than benthic lineages (Sims et al. 2006), and freshwater lineages are generally nested within, and younger than their marine counterparts (Alverson 2014; Ruck et al. 2016). A superficial look at these data might suggest that younger lineages are disproportionately more diverse than older ones, perhaps due to differences in diversification dynamics. We compared a total of 23 models (Table S1): 11 in the base set comparing character-dependent to character-independent models, six testing specific hypotheses related to diversification parameters, and another six comparing hypotheses about transition rates. From these 23 models, four were supported by an Akaike weight (**ω** ≥ 1%; Table 4) and were retained for further analyses using model averaging (Table 4). These included a model where the plankton-benthos contrast had no effect (MP = MB and FP = FB, **ω** = 84%), the most complex MuHiSSE model without constraints on turnover (**ω** = 10%), a model that constrained the transition rates such that the marine plankton was absorbing (MP -> MB = 0, but MP -> FP *Φ* 0, ≠ = 5%), and a model where the freshwater plankton was absorbing (FP -> MP = 0, but FP -> FB ≠ 0, **ω** = 1%).

**Table 4.** Comparison of character-dependent and independent models for the interaction between diversification and the combined salinity+habitat states. Shown are the models with support ≥ 1% used in model-averaging. For the full set of results refer to Supplementary table S1 Abbreviations: npd, number of diversification parameters (turnover and extinction fraction); npt, number of transition parameters; Δ, difference in AlCc; ω, Akaike weight; *ε*, extinction fraction.

| Model | Parameter setup | lnL | npd | npt | AICc | $\Delta$ | $\omega$ |
| --- | --- | --- | --- | --- | --- | --- | --- |
| No habitat effect | $\epsilon$=fix; MP = MB and FP = FB | -5518.47 | 7 | 18 | 11083.93 | 0 | 0.84 |
| MuHiSSE full | $\epsilon$=fix; no additional constraints | -5516.44 | 9 | 18 | 11088.25 | 4.32 | 0.10 |
| MP absorbing plankton | $\epsilon$=fix; MP → MB = 0, but MP → FP ≠ 0 | -5519.25 | 9 | 17 | 11089.68 | 5.75 | 0.05 |
| FP absorbing freshwater | $\epsilon$=fix; FP → MP = 0, but FP → FB ≠ 0 | -5520.47 | 9 | 17 | 11092.12 | 8.19 | 0.01 |

Net turnover and net diversification in freshwater habitats were 3 times faster than in marine environments after averaging across these four models and over their hidden-state ancestral reconstructions (Fig. 1C). Lineages with suspended and substrate-associated growth habits had similar turnover rates in both marine and freshwaters due to the weight of the model without a plankton-benthos effect. The waiting times to a net gain of one species estimated here were in general agreement with previous estimates of net diversification in diatoms (Nakov et al. 2018). For example, the net diversification waiting time for marine planktonic lineages under the MuHiSSE model was 17 My in one and 2.7 My in the second hidden state, whereas the waiting time for oogamous nonmotile diatoms (most common in the marine plankton) was previously estimated as between 16.6 and 100 My (phylogenetic data) or roughly 34.5 My (fossil data) (Nakov et al. 2018). The relatively high global extinction fraction (ε = 0.8; Table S2), estimated here from extant species only, was similar to calculations made from Cenozoic marine fossils (ε = 0.75; Nakov et al. 2018). Analyses using alternative implementations of the conditioning of survival of crown lineages indicated that the two approaches produce very similar results (Fig. S1, comparison for the base models).

### Taxonomic bias

Like any diversification analysis that accounts for unsampled diversity, our results rely on the assumption that the total (described and undescribed) species richness in our focal regimes is directly proportional to the diversity described from those habitats to date. However, in addition to differences in diversification rates, these lopsided diversity ratios could also arise from unequal taxonomic effort or from nonstandard taxonomic practices (e.g., splitting vs. lumping) between researchers studying different habitats. It is difficult to assess how these problems might affect our analyses, or analyses of any group, without some non-arbitrary method of scaling the diversity data or sampling fractions to account for possible unknown biases related to, for example, over-splitting (e.g., in freshwater taxa) or lower taxonomic research effort (e.g., in marine taxa). Nevertheless, we repeated the marine-freshwater analysis with arbitrarily reduced marine sampling fractions, from the observed value of 21% down to 14%, 10%, 6%, and 2%. Irrespective of how low we set the marine sampling fraction, we found that HiSSE models were still favored over character-independent models. With smaller sampling fractions, the diversification and turnover rate estimates for marine lineages increased, but the signal for faster diversification in freshwaters, at least in one of the hidden states, was detected in all but the analysis with a marine sampling fraction of 2%. In the latter, marine lineages diversified faster than freshwater in both hidden states. Overall, these results indicated that, if the past nearly 250 years of taxonomic work on diatoms is completely biased and non-representative of diatom diversity as a whole, then some of our inferences might be affected.

To evaluate potential clade-specific sampling biases, we repeated the analysis of the entire diatom phylogeny on four subclades with sufficient sampling across the two environmental contrasts. Although it is not expected that model selection or parameter estimates from the entire phylogeny should match those inferred from subtrees that differ, among others, in age, diversity, ecology, life history, and genomic attributes (Beaulieu and O’Meara 2018; Rabosky 2019), we found that the global result, i.e., faster turnover in freshwater compared to marine lineages, was qualitatively replicated in most of the subtrees (see Supplementary Text 1 and Fig. S2 for details).

## Discussion

One of the most prominent features of aquatic biodiversity is the disparity in species richness between marine and freshwater habitats. Marine habitats are geologically old, have existed continuously since the formation of the oceans and amount to >99% of the habitable water on Earth, whereas present-day freshwater habitats are much younger and have shorter lifespans determined by glaciation, droughts, or tectonic events. Surprisingly, in lineages with substantial diversity across the salinity gradient, the younger and derived freshwater clades are often more species rich than their marine counterparts. This difference in diversity is generally a consequence of higher diversification rates in freshwater environments relative to marine habitats. However, freshwater lineages also tend to exhibit higher turnover rates, which might partly explain the preponderance of diverse clades nearer the present. Pulses of speciation followed by pulses of extinction are indicative of higher turnover rates. Importantly, differences in turnover rates suggest that transitions into and out of freshwater environments jointly impact speciation and extinction rates.

### Rate variation and hidden states

Our analyses of the marine-freshwater and plankton-benthos contrasts, considered either independently or in combination, revealed strong support for hidden state models, either character-dependent (salinity and combined) or character-independent (habitat). This indicates, as would be expected for a lineage as phylogenetically and ecologically diverse as diatoms, that the focal characters alone (salinity, habitat or their combination) do not explain all of the variation in diversification rates across the phylogeny, even after partitioning the data into four regimes corresponding to the states of the combined habitat+salinity character. This further underscores the need to account for the multitude of environmental, ecological, and trait factors that were not explicitly considered here, but whose cumulative effect on diversification might be comparable to the effects of salinity and habitat. To illustrate this point, we can compare the differences in diversification between environments to the differences between hidden states within an environment. Marine planktonic lineages reconstructed in one of the hidden states had a waiting time to a speciation event of about 17 My. This was similar to marine benthic lineages in the same hidden state (17 My), but was several times longer than marine planktonic lineages in the other hidden state (2.7 My; Fig. 2; Table S2). In other words, growing suspended in the water column or associated with a substrate conferred a smaller difference in diversification than differences conferred by unobserved factors approximated by the hidden states (Fig. 2; Table S2). This is not to say that salinity has not played an important role in diatom diversification, but the opposite—that its effect is strong enough to be detected despite broad variation related to other, unobserved factors.

**Figure 2.**
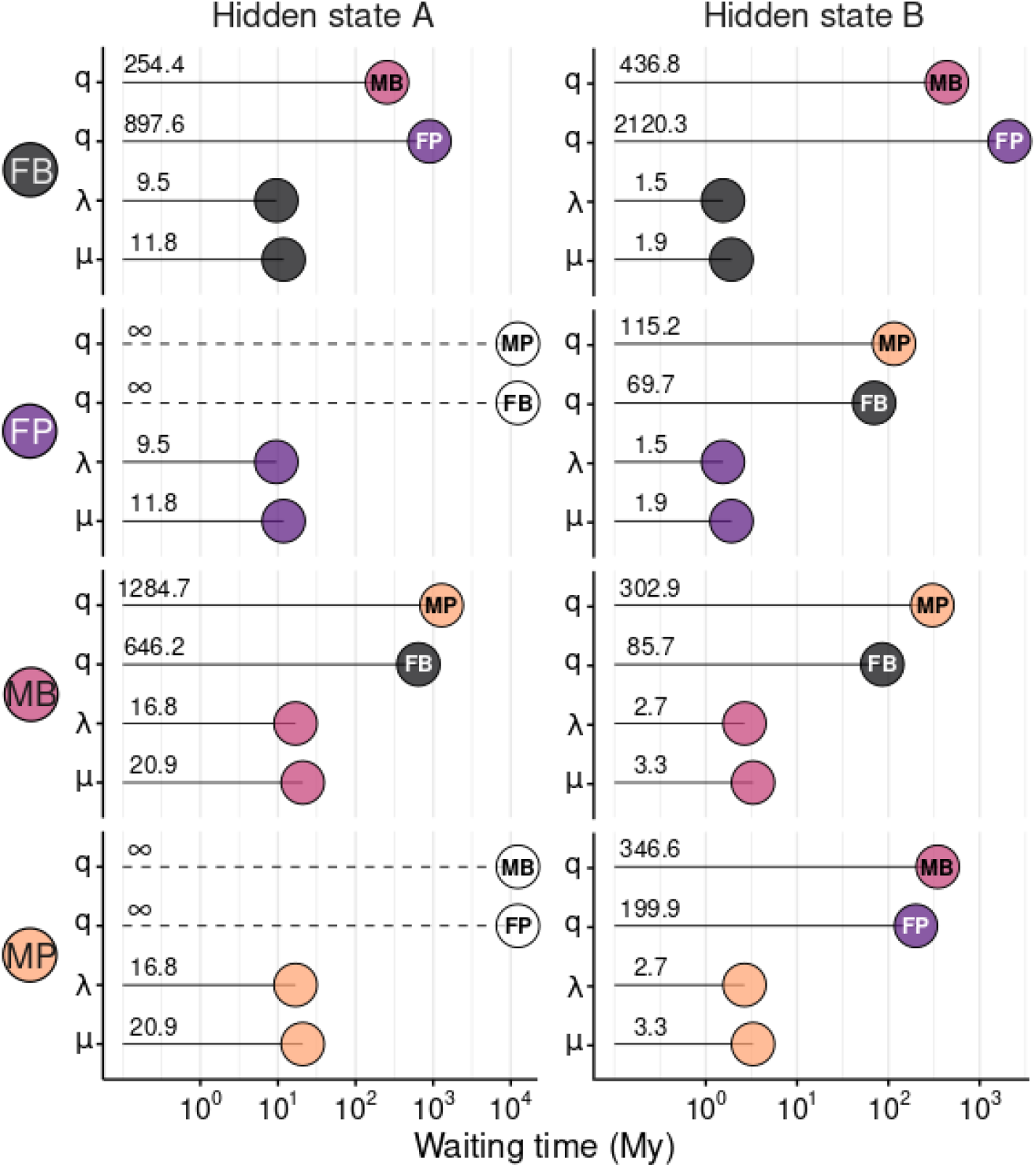
Waiting times (inverse of the rates) in million years (My) for speciation, extinction, and transitions for the habitat + salinity under the favored MuHiSSE model (Table 4). The source environment is denoted with a two-letter abbreviation on the y axis. For speciation (λ) and extinction (μ) the color of the end-point circle matches the source environment. For transition rates (q), the color and abbreviation at the end-point circle denote the destination of colonization. Numbers along the lines are the estimated waiting times. Transitions rates estimated as indistinguishable from zero (and, consequently, infinity waiting times) are shown with dashed lines and open circles. Left and right side panels correspond to the two hidden states. Abbreviations: FB, freshwater--benthos; FP, freshwater--plankton; MB, marine--benthos; MP, marine--plankton.

It is worth considering what some of these unmeasured factors might be. For example, differences between lotic and lentic habitats, and within the latter, differences in age, size, depth, and longevity among lakes of glacial, tectonic, and volcanic origin might be driving some of the variation within freshwaters. Likewise, the drastic contrast in temperature, nutrients, and light conditions experienced by diatom communities in coastal areas vs. nutrient-rich marine polar waters or oligotrophic oceanic gyres could account for some of the hidden state components within marine lineages. Water temperature in particular appears to be an important but untested covariate, as it changes with latitude, geography (coastal vs. pelagic), and water depth (linked to our plankton-benthos contrast), and is related to metabolic rates that one could easily envision influencing diversification dynamics. For example, Crampton et al. (2016) found that peaks in species turnover of marine Antarctic diatoms coincided with cooling of southern latitudes and expansion of ground and sea ice. Five such pulses above the baseline turnover have been inferred over the past 15 My (Crampton et al. 2016), contributing to relatively fast turnover within Antarctic diatom communities. Similar cooling-driven fluctuations in turnover are less likely to have occurred in tropical ocean regions where temperature has been relatively constant over the past 15 My (Powell and Glazier 2017). Therefore, it is possible that a portion of the variation captured by the hidden states might also be related to temperature and latitude. In this regard, diatoms are among the few lineages with a reverse latitudinal diversity gradient, which is thought to be a product of asymmetric range expansion toward higher latitudes (Powell and Glazier 2017).

Variation in rates of diversification within regimes might also stem from other traits that covary with diversity patterns in diatoms. For example, motility of vegetative cells, which allowed colonization of new types of habitats and fundamentally changed how these diatoms interact with each other and the environment, was also associated with a faster rate of diversification (Nakov et al. 2018). A comparison between motile and non-motile lineages showed that the probability of occupying the second hidden state—where net turnover was much faster (Fig. 3, Table S2)—was higher in motile lineages, though this result was not significant (phylogenetic ANOVA, p=0.4). It is important to reiterate that this *post hoc* exploration of potential hidden variables is not meant to draw conclusions about other factors not directly tested here, but only as an illustration of the many other variables that may have interacted with diatom diversification (e.g., Lewitus et al. 2018).

**Figure 3.**
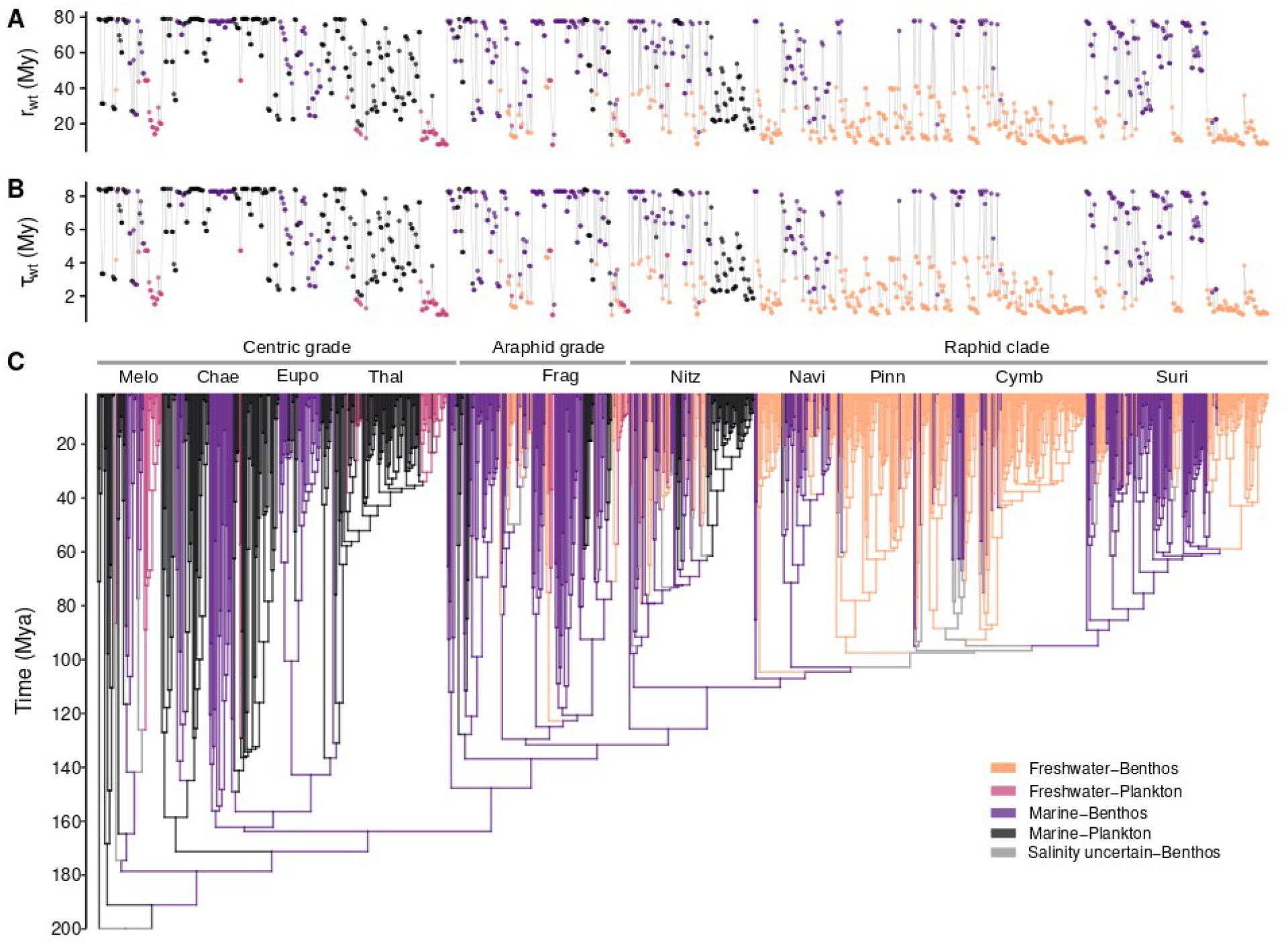
Ancestral state reconstructions of habitats and tip estimates of net turnover (*τ*, speciation + extinction) and net diversification (*r*, speciation - extinction) rates inverted to waiting times (wt) in millions of years. The high-turnover and high-diversification freshwater lineages (shorter waiting times) are predominantly benthic within the raphid clade with motile vegetative cells (e.g., Suri and Cymb) and predominantly planktonic within the non-motile centric and araphid grades (e.g., Frag, Thal, and Melo). Abbreviations: Mya, millions of years ago; Melo, Melosirales; Chae, Chaetocerotales; Eupo, Eupodiscales; Thal, Thalassiosirales; Frag, fragilaroids; Nitz, Bacillariales; Navi, naviculoids; Pinn, pinnulariids; Cymb, Cymbellales; Suri, Surirellales; Parm, Parmales; Lept, Leptocylindrales + Corethrales; Cosc, Coscinodiscales + Rhizosoleniales.

### Diatom ancestry, absorbing habitats, and phylogenetic diversity

Ancestral state reconstructions supported a marine planktonic environment for the most recent common ancestor of diatoms, but they quickly colonized the marine benthos (see also Kooistra et al. 2007). The oldest extant benthic lineages were estimated to be about 180-175 My old (Fig. 3). Colonizations of freshwater did not occur until much later—the oldest extant lineages to gain a foothold in the freshwater plankton date back to 120-130 Mya, some 60 My after the origin of diatoms (Fig. 3). Apart from a few isolated cases that have not left a substantial mark on present day diversity (e.g., *Ellerbeckia, Terpsinoe*, and *Pleurosira)*, diatoms did not diversify greatly within the freshwater benthos until after the evolution of anisogamous sexual reproduction and axial cell symmetry in the pennate diatom lineage (Fig. 3). Marine benthic ancestry was inferred for the MRCA of the pennate clade, and within it, the clade of actively motile raphid diatoms (Fig. 3). Overall, although our reconstructions supported a planktonic ancestry for diatoms, it is clear that the marine benthos has been the center stage of diatom evolution, seeding diversity across a range of environments (e.g., the marine plankton and freshwater benthos) throughout their history (Fig 3; see also Kooistra et al. 2007).

The permeability of the salinity barrier is a recurring topic in diatom ecology and evolution. Historically, both in diatoms, and also more broadly across protists (Mann 1999; Cavalier-Smith 2009; Logares et al. 2009), marine-to-freshwater colonizations have been thought of as difficult and irreversible transitions, which led to the development of the ‘Rubicon’ hypothesis, which states that once a lineage colonizes freshwaters there is no return to the marine environment (Mann 1999). By simultaneously estimating diversification and transition rates, our models provided a powerful test of this hypothesis, and by accounting for rate heterogeneity within regimes we could test whether salinity is a Rubicon in all, or just some, diatom lineages. Models that disallowed freshwater-to-marine or plankton-to-benthos transitions as required to directly test these hypotheses received no support (Table 1, 4). However, our analyses showed that the rates of traversal of either environmental contrasts were dependent on hidden factors. For example, in some lineages, freshwater-to-marine colonizations were rare and marine-to-freshwater far more common, whereas in others the waiting time to recolonization of marine habitats was 13 times shorter than the waiting time of marine-to-freshwater transitions (Table 2). Models that accounted for the possibility that marine-to-freshwater transitions differ between planktonic and benthic species, or that plankton-to-benthos migration differs between marine and freshwater lineages, showed similar variation (MuHiSSE analysis, Fig. 2, Table S2). Therefore, although the notion of irreversibility of freshwater or plankton colonizations is not supported for the diatom lineage as a whole, it nonetheless might apply to a subset of diatom lineages in which recolonization of ancestral habitats, although still possible, takes far longer (Fig. 2, Table 2, 3).

Comparison of the transition rates can also be informative about the phylogenetic diversity of communities occupying the extremes of these environmental contrasts. This can be illustrated by comparing the rates of outflow from the freshwater plankton between the first and second hidden state. For example, in some lineages escaping the freshwater plankton was a rare event, but in others, outflow from the freshwater plankton was among the most common transitions with the shortest retention time (Fig. 2). Such contrasts in transition rates suggest that for some lineages the freshwater plankton is a transient state, which, combined with the generally short-lived nature of freshwater planktonic habitats over deep geologic timescales, suggests an explanation regarding the generally narrow phylogenetic breadth of freshwater planktonic diatom communities. The few lineages that have diversified substantially in the freshwater plankton are almost entirely restricted to this habitat and have mostly been sourced from the marine plankton (e.g., the genus *Aulacoseira* within marine planktonic *Melosirales* or cyclostephanoid diatoms within the ancestrally marine planktonic *Thalassiosirales).* In turn, for most of the remaining lineages, colonizations of freshwater plankton were sporadic and did not lead to substantial species diversification (Fig. 3). This is in contrast to the much more phylogenetically diverse marine plankton, which has been colonized repeatedly by many marine benthic lineages (Fig 3; see also Kooistra et al. 2007), harbors a considerable diversity of araphid (e.g., *Asterionellopsis* and *Thalassiosinema)* and raphid diatoms (e.g., *Fragilariopsis* and *Pseudo-nitzschia;* Fig. 3), and as suggested by the rates of outflow from the freshwater plankton, is also recolonized by freshwater planktonic lineages (Fig. 2, Table S2). These observations, combined with the non-uniform recruitment to the freshwater plankton—several old and species-rich marine planktonic lineages have not crossed into freshwaters or remain disproportionately more diverse in saline waters (e.g., Rhizosoleniales, Coscinodiscales, and Chaetocerales; Fig. 3)—also suggest that marine planktonic diatom assemblages are phylogenetically and functionally more diverse than their freshwater counterparts.

It is important to consider here that transitions between marine and freshwaters, or plankton and benthos, certainly do not occur instantaneously. Rather, the most likely colonizers of freshwater are marine lineages already capable of growing at a lower salinity (brackish) or a broader range of salinity (euryhaline), for example species living in estuaries or saline inland lakes. A more appropriate model for these analyses would therefore account for the presence of euryhaline or brackish species and potentially constrain the transitions between the salinity extremes by forcing an intermediate euryhaline or brackish state. In fact, an analysis of a small clade of predominantly benthic diatoms did support a model of ordered, reversible marine<->euryhaline<->freshwater transitions, although, it did not account for diversification (Ruck et al. 2016). Replicating such an analysis at the level of all diatoms and in a SSE modeling framework would certainly be a welcome approach. Using geographic SSE models (GeoSSE, Goldberg et al. 2011; or HiGeoSSE, Caetano et al. 2018) where species with euryhaline ranges diverge into freshwater and marine populations is also an interesting way to model these data (see e.g., Betancur-R et al. 2015). However, implementation of such approaches is hampered at this time due to the lack of data for the range of salinity tolerance across the diatom tree, the patchy information on diatoms in biogeographic databases, and the resulting uncertainties in estimation of total diversity and sampling fractions for euryhaline or brackish species.

In sum, the marine-freshwater and plankton-benthos contrasts are unquestionably important determinants of aquatic diversity. In diatoms, a species-rich lineage of algae important for global ecology and biogeochemistry, we found that the marine-freshwater diversity imbalance is maintained by faster turnover and faster net diversification in freshwaters. This pattern holds for both planktonic and benthic habitats, an environmental divide that itself has not greatly impacted diatom diversification. More generally, our analyses showed that global diversity patterns are unlikely to be attributable to a few, broadly defined regimes. Rather they are the product of many potentially interacting traits and environmental factors whose individual or aggregate effects might equal or surpass the influence of factors traditionally considered important. Our analyses help illustrate how consideration of multiple possibly interacting characters provides a more refined understanding of diversification in old, widespread, and species-rich clades.

## Supplementary information

### Clade-specific sampling biases

Clade-specific sampling biases could arise as a result of deeper phylogenetic sampling in lineages of ecological, biotechnological, or public health interest or due to natural variation in the distribution of lineages across the focal environmental contrasts. One way to assess potential biases of this type is to check whether a pattern observed across the entire phylogeny is replicated at shallower phylogenetic scales. There is no reason to expect that the same models should be favored or similar parameters should apply for subtrees that differ substantially in age, ecology, life history, genomic features, and so on (Beaulieu and O’Meara 2018; Rabosky 2019). But it might be informative to evaluate in broad terms whether a global result, i.e., turnover in freshwater lineages is faster than in marine, is detectable, at least to some degree, in younger, smaller, clades.

To address this, we analyzed four subtrees with sufficient representation across both the marine-freshwater and plankton-benthos contrasts (Fig. S2) and asked whether the model favored for the all-diatom tree was also favored in the subtrees and whether model-averaged tip-associated turnover rates estimated for the subtrees recovered a pattern similar to that of the all-diatom phylogeny. For these smaller clades, the CID4 model was favored over HiSSE in all but one clade, for both salinity and habitat. However, even though a model of character-dependent diversification was not universally supported, model-averaged tip rates showed that, on average, freshwater species had higher turnover than marine in all but one clade. For habitat, in some clades turnover in the plankton was faster, on average, than in the benthos, but in others, the two habitats had essentially the same mean rates, consistent with the pattern observed at the all-diatom scale. For the two largest clades, raphid and multipolar diatoms, we also repeated the combined habitat+salinity analysis and found that model selection was replicated for the clade of raphid diatoms but not for the clade of multipolar lineages. Model-averaged rates again resembled the pattern observed for the whole diatom tree, with turnover faster in freshwater than marine environments.

**Table S1.** The full set of multistate models examined for the combined salinity (marine-freshwater) + habitat (plankton-benthos) character. In all cases the extinction fraction (*ε*, speciation/extinction) was held constant across states, while turnover (*τ*, speciation + extinction) was free to vary across states or constrained to test specific hypotheses.

| Model | Parameter setup | lnL | AICc | dAIC | AICw |
| --- | --- | --- | --- | --- | --- |
| No habitat effect | MP = MB, FP = FB | -5518.47 | 11083.93 | 0 | 0.84 |
| MuHiSSE relax | No additional constraints | -5516.44 | 11088.25 | 4.32 | 0.1 |
| MP absorbing plankton | MP → MB = 0, MP → FP ≠ 0 | -5519.25 | 11089.68 | 5.75 | 0.05 |
| FP absorbing freshwater | FP → MP = 0, FP → FB ≠ 0 | -5520.47 | 11092.12 | 8.19 | 0.01 |
| No salinity effect in plankton | MP = FP, MB ≠ FB | -5524.71 | 11100.6 | 16.67 | 0 |
| FP absorbing plankton | FP → MP ≠ 0, FP → FB = 0 | -5527.68 | 11106.53 | 22.6 | 0 |
| No habitat effect in freshwater | MP ≠ MB, FP = FB | -5530.05 | 11111.28 | 27.35 | 0 |
| No habitat effect in marine | MP = MB, FP ≠ FB | -5530.4 | 11111.98 | 28.05 | 0 |
| No salinity effect in benthos | MP ≠ FP, MB = FB | -5533.29 | 11117.75 | 33.82 | 0 |
| No salinity effect | MP = FP, MB = FB | -5536.56 | 11120.11 | 36.18 | 0 |
| MuHiSSE | Symmetric transitions between hidden states | -5539.58 | 11132.43 | 48.5 | 0 |
| Plankton absorbing | FP → FB = 0, MP → MB = 0 | -5542.91 | 11132.81 | 48.88 | 0 |
| CID4 relax | No additional constraints | -5563.63 | 11161.81 | 77.88 | 0 |
| CID5 relax | No additional constraints | -5562.7 | 11164.08 | 80.15 | 0 |
| CID3 relax | No additional constraints | -5566.86 | 11164.15 | 80.22 | 0 |
| CID6 relax | No additional constraints | -5562.69 | 11168.22 | 84.29 | 0 |
| CID2 relax | No additional constraints | -5572.27 | 11170.86 | 86.93 | 0 |
| FB absorbing freshwater | FB → MB = 0, FB → FP ≠ 0 | -5560.22 | 11171.61 | 87.68 | 0 |
| CID7 relax | No additional constraints | -5562.68 | 11172.36 | 88.43 | 0 |
| CID8 relax | No additional constraints | -5562.63 | 11176.43 | 92.5 | 0 |
| Freshwater absorbing | FB → MB = 0, FP → MP = 0 | -5594.11 | 11235.21 | 151.28 | 0 |
| MuSSE | No additional constraints | -5701.85 | 11430.03 | 346.1 | 0 |
| Dull | No additional constraints | -5903.94 | 11828.08 | 744.15 | 0 |

**Table S2.**
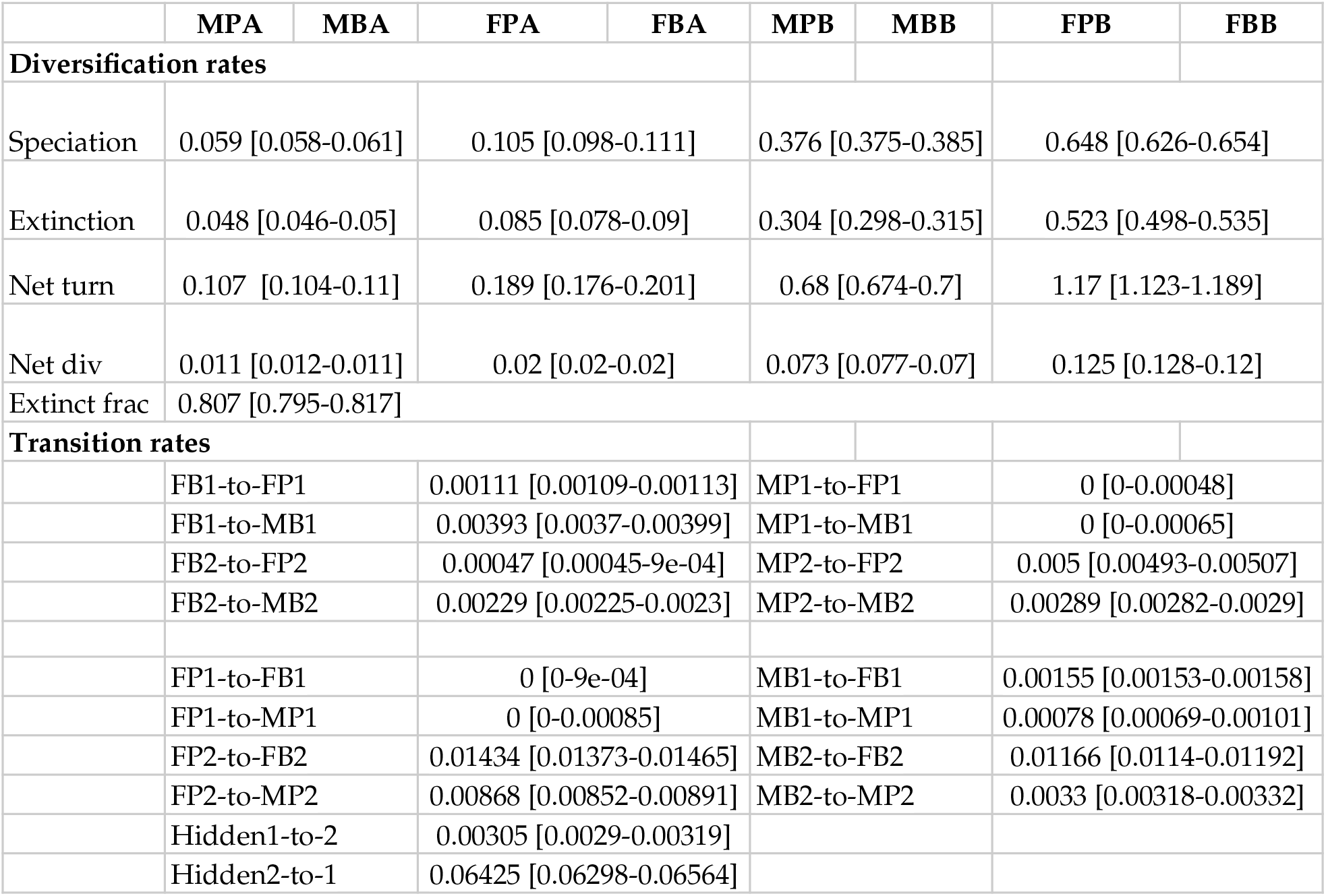
Parameter estimates and 2 log likelihood support region (in brackets) for the combined salinity + habitat character states [marine-plankton (MP), marine-benthos (MB), freshwater-plankton (FP) and freshwater-benthos (FB)]. The favored model (*ω* = 0.84) was hidden state multistate speciation and extinction without a habitat effect on diversification (Table 4). The letters next to character states indicate the hidden states.

|  | MPA | MBA | FPA | FBA | MPB | MBB | FPB | FBB |
| --- | --- | --- | --- | --- | --- | --- | --- | --- |
| Diversification rates |  |  |  |  |  |  |  |  |
| Speciation | 0.059 [0.058-0.061] |  | 0.105 [0.098-0.111] |  | 0.376 [0.375-0.385] |  | 0.648 [0.626-0.654] |  |
| Extinction | 0.048 [0.046-0.05] |  | 0.085 [0.078-0.09] |  | 0.304 [0.298-0.315] |  | 0.523 [0.498-0.535] |  |
| Net turn | 0.107 [0.104-0.11] |  | 0.189 [0.176-0.201] |  | 0.68 [0.674-0.7] |  | 1.17 [1.123-1.189] |  |
| Net div | 0.011 [0.012-0.011] |  | 0.02 [0.02-0.02] |  | 0.073 [0.077-0.07] |  | 0.125 [0.128-0.12] |  |
| Extinct frac | 0.807 [0.795-0.817] |  |  |  |  |  |  |  |
| Transition rates |  |  |  |  |  |  |  |  |
|  | FB1-to-FP1 |  | 0.00111 [0.00109-0.00113] |  | MP1-to-FP1 |  | 0 [0-0.00048] |  |
|  | FB1-to-MB1 |  | 0.00393 [0.0037-0.00399] |  | MP1-to-MB1 |  | 0 [0-0.00065] |  |
|  | FB2-to-FP2 |  | 0.00047 [0.00045-9e-04] |  | MP2-to-FP2 |  | 0.005 [0.00493-0.00507] |  |
|  | FB2-to-MB2 |  | 0.00229 [0.00225-0.0023] |  | MP2-to-MB2 |  | 0.00289 [0.00282-0.0029] |  |
|  | FP1-to-FB1 |  | 0 [0-9e-04] |  | MB1-to-FB1 |  | 0.00155 [0.00153-0.00158] |  |
|  | FP1-to-MP1 |  | 0 [0-0.00085] |  | MB1-to-MP1 |  | 0.00078 [0.00069-0.00101] |  |
|  | FP2-to-FB2 |  | 0.01434 [0.01373-0.01465] |  | MB2-to-FB2 |  | 0.01166 [0.0114-0.01192] |  |
|  | FP2-to-MP2 |  | 0.00868 [0.00852-0.00891] |  | MB2-to-MP2 |  | 0.0033 [0.00318-0.00332] |  |
|  | Hidden1-to-2 |  | 0.00305 [0.0029-0.00319] |  |  |  |  |  |
|  | Hidden2-to-1 |  | 0.06425 [0.06298-0.06564] |  |  |  |  |  |

**Figure S1.**
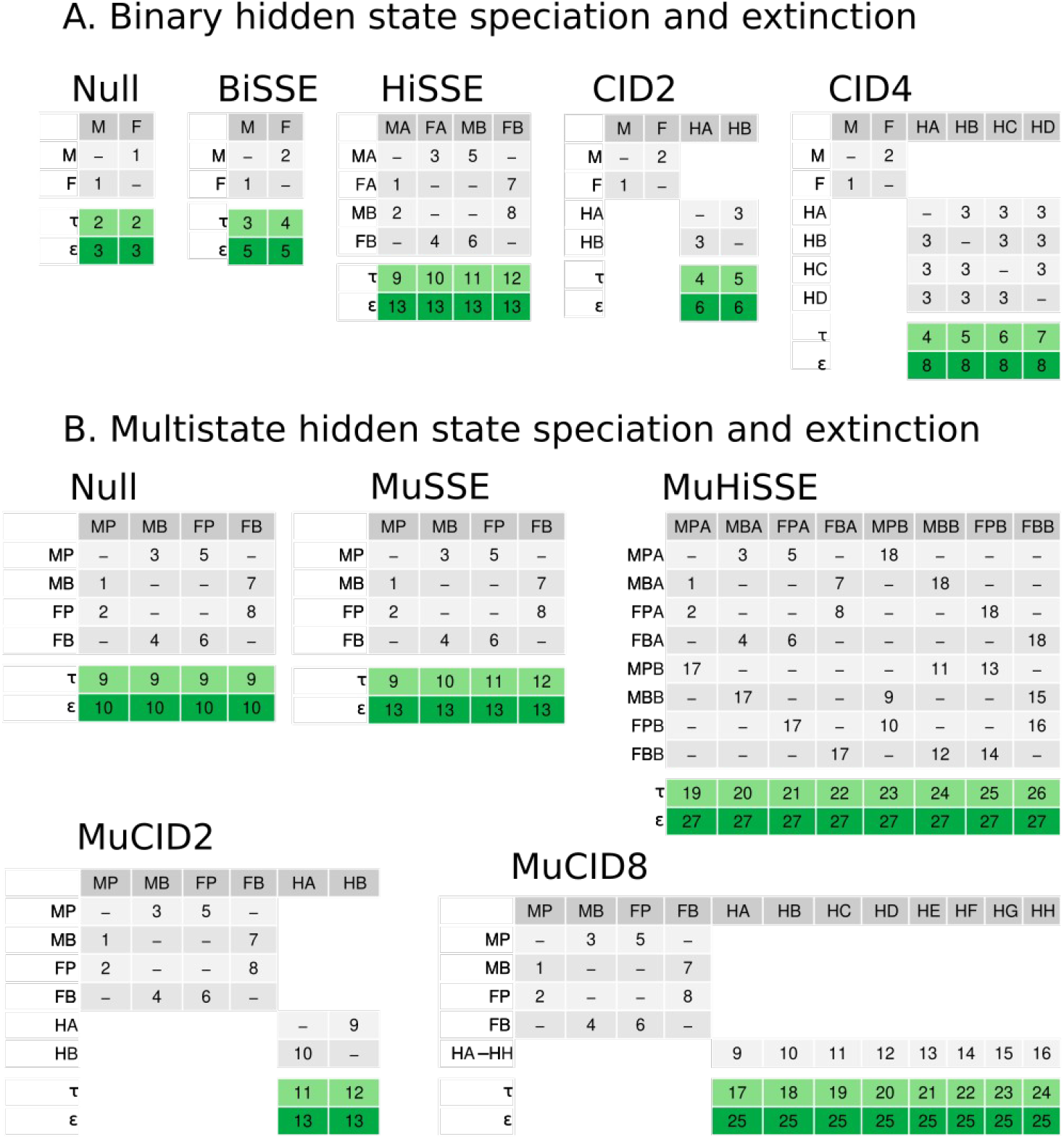
Character-dependent and character-independent diversification models for binary (A) and multistate (B) coding of salinity and habitat. The top part (grey) depicts the transition rate matrix with row and column names representing observed character states (M=marine, F=freshwater and their combinations) or hidden (H) states. The letters A-H (e.g., MA, MB, FA, FB) differentiate hidden states within a character state. The bottom part (green), labeled with τ and ε shows the parameters for net turnover (speciation+extinction) and extinction fraction (extinction/speciation), respectively. Each τ and ε parameter is linked to the state denoted in the column name. Note that diversification parameters are associated to character states in SSE models, but not in CID models, where they are linked to hidden states. Numbers inside the table represent the indexes of model parameters. For constrained parameters, the same index occurs in multiple cells (e.g., ε was fixed in all models, so it has one parameter in each table). Models for habitat (plankton-benthos) were the same as the binary models for salinity (A).

**Figure S2.**
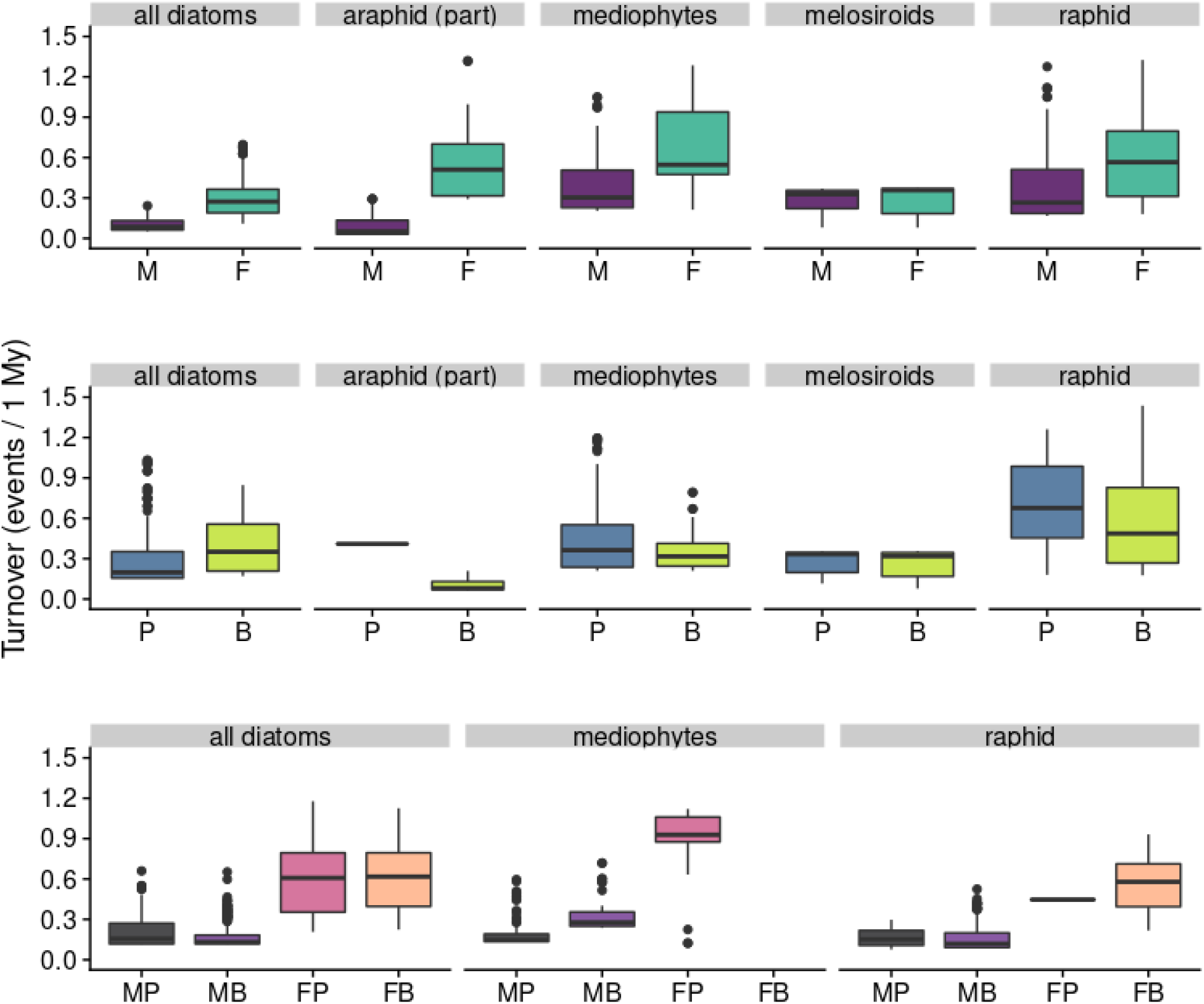
Tip estimates of net turnover (speciation + extinction) averaged across models and hidden states reconstructions. Shown are the clades of that had at least 40 sampled species and reasonable representation in marine and freshwaters, as well as plankton and benthos. In addition to the entire diatoms phylogeny (all diatoms), these clades include the raphid diatoms (raphid), part of araphid diatoms (araphid part), melosiroid diatoms (melosiroid), and multi-polar diatoms (mediophytes). Top row: binary coding for marine vs. freshwater; Middle row: binary coding for planktonic vs. benthic; Bottom row: multi-state coding for combined habitat+salinity states. Abbreviations: M, marine; F, freshwater; P, plankton; B, benthos.

**Figure S3.**
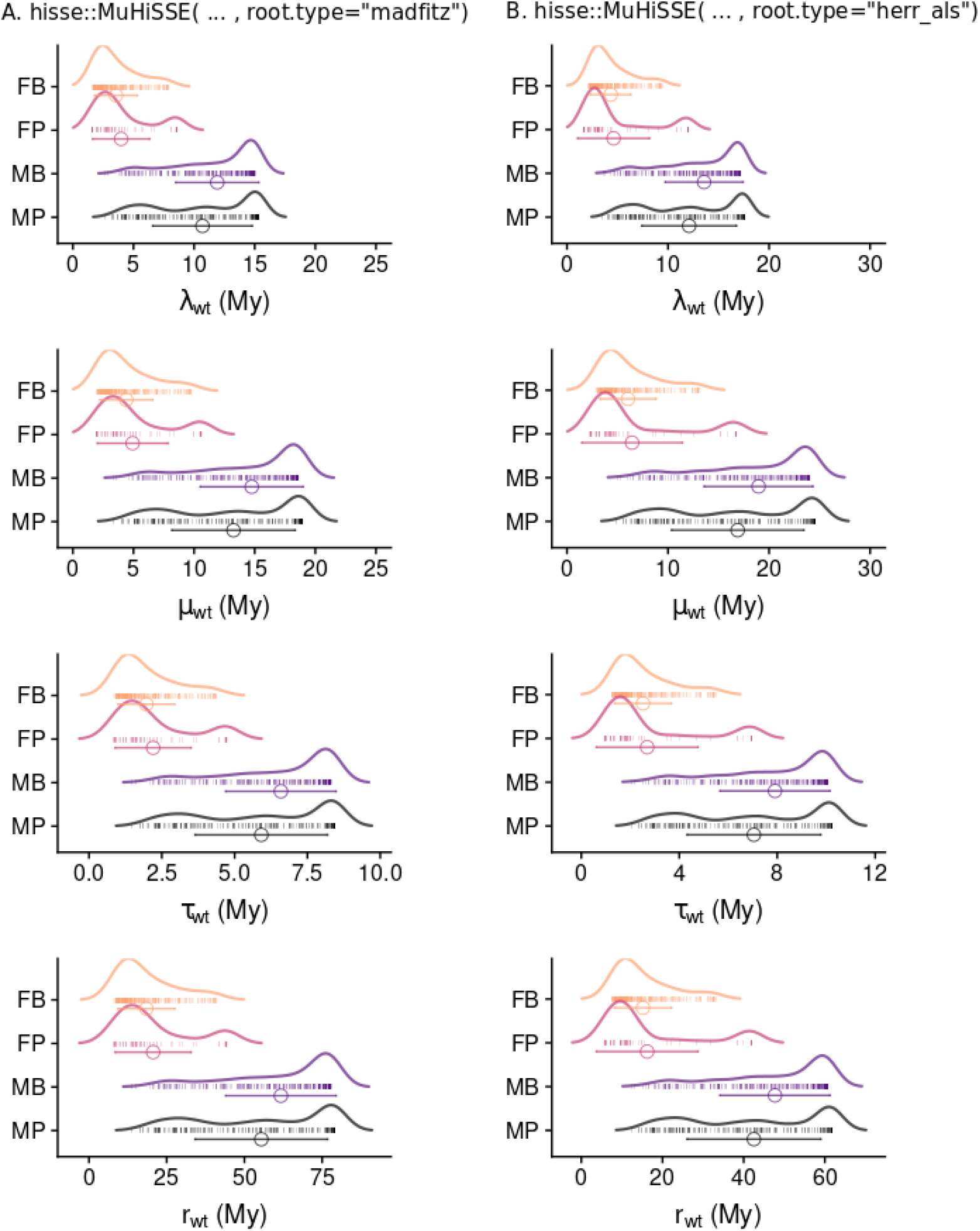
Distributions of tip-estimates of speciation (λ), extinction (μ), net turnover (*τ*, speciation + extinction) and net diversification (***r***, speciation - extinction) averaged across hidden states under the MuHiSSE model and two alternative implementations of the conditioning for survival of crown lineages (FitzJohn et al. 2009; Herrera-Alsina et al. 2018). The circles and horizontal error bars show the mean ± standard deviation across tips of the phylogeny and the carpet points show the individual rates. The rates are inverted into waiting times (wt) in millions of years. Abbreviations: M, marine; F, freshwater; P, plankton; B, benthos.

**Figure S4.**
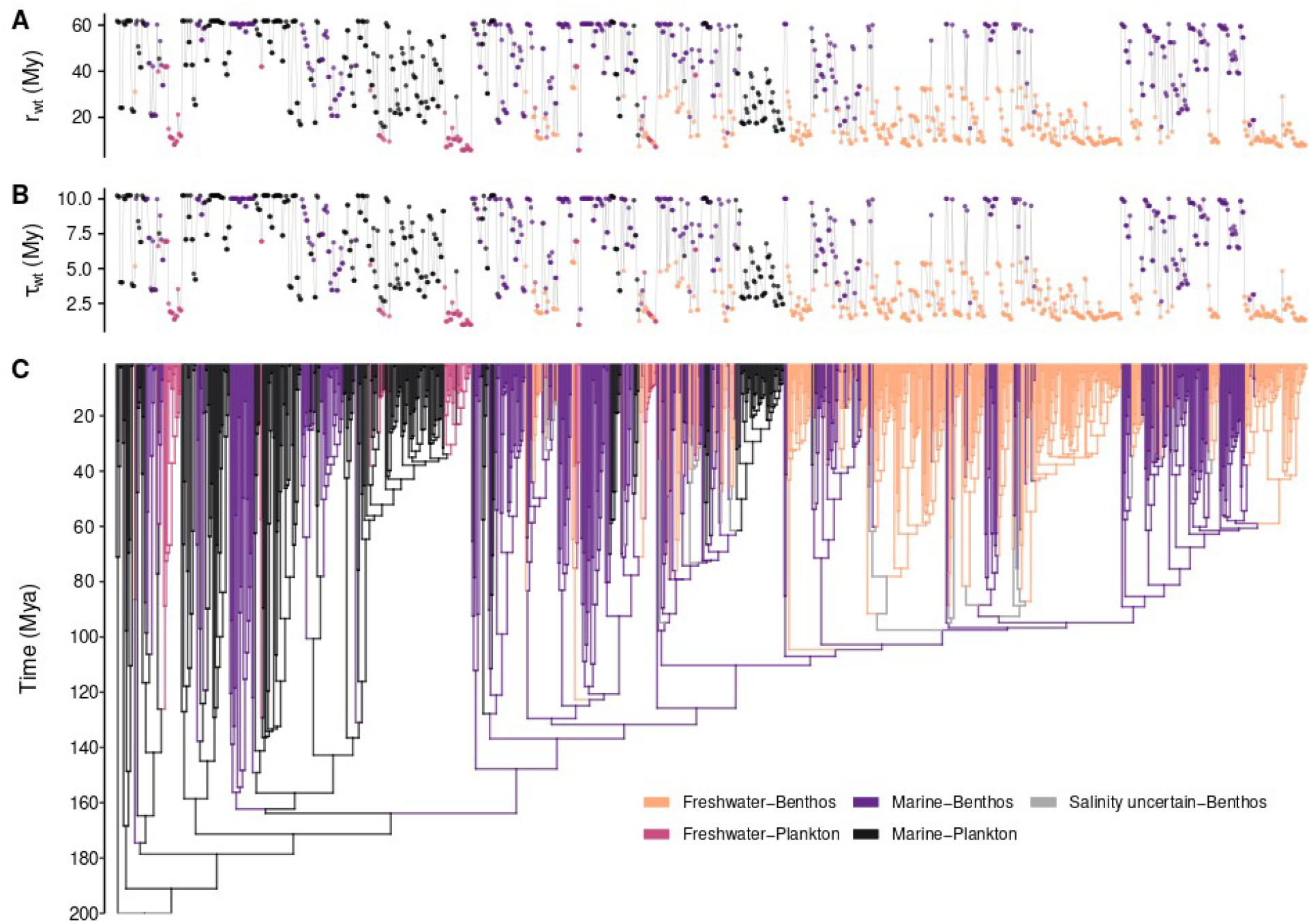
Ancestral state reconstructions of environments and the tip estimates of net turnover (*τ*, speciation + extinction) and net diversification (***r***, speciation - extinction) following Herrera-Alsina et al.’s **(2019)** implementation of the conditioning for survival of crown lineages. Rates were inverted to waiting times (wt) in millions of years. The high-turnover and high-diversification freshwater lineages (lower waiting times) are predominantly benthic among motile raphid pennate diatoms (e.g., Suri and Cymb) and predominantly planktonic within the non-motile centric and araphid grades (e.g., Frag, Thal, and Melo). Abbreviations: Mya, millions of years ago; Melo, Melosirales; Chae, Chaetocerotales; Eupo, Eupodiscales; Thal, Thalassiosirales; Frag, fragilaroids; Nitz, Bacillariales; Navi, naviculoids; Pinn, pinnulariids; Cymb, Cymbellales; Suri, Surirellales; Parm, Parmales; Lept, Leptocylindrales + Corethrales; Cosc, Coscinodiscales + Rhizosoleniales.

